# Lineage-specific LTR Retrotransposon Repertoires Engage OCT4–SOX2–SMARCA4 Chromatin Regulatory Architecture in Mouse and Human Pluripotent Stem Cells

**DOI:** 10.64898/2026.11.23.768445

**Authors:** Hikaru Suzuki, Kenji Ichiyanagi

## Abstract

**Background:** Long terminal repeat (LTR) retrotransposons contribute to regulatory sequences in mammalian genomes and provide binding sites for pluripotency-related transcription factors (TFs), such as POU5F1 (also called OCT4). SMARCA4 (also called BRG1), the ATPase catalytic subunit of BRG1/BRM-associated factor (BAF) chromatin-remodeling complexes, cooperates with pluripotency-related TFs and regulates chromatin accessibility. However, whether SMARCA4 preferentially occupies specific LTR subfamilies, whether it promotes chromatin accessibility at these elements, and how its occupancy relates to that of pluripotency-related TFs remains unclear. Here, we investigated the relationship between LTR retrotransposons and POU5F1–SOX2–SMARCA4 chromatin regulatory mechanisms in mouse and human pluripotent stem cells.

**Results:** Analysis of publicly available datasets revealed that SMARCA4 preferentially occupies specific LTR subfamilies, including RLTR13D6 and RLTR9E, in mouse embryonic stem cells. These LTRs exhibited accessible chromatin and active histone modifications, and were co-occupied by POU5F1 and SOX2. SMARCA4 depletion was associated with reduced chromatin accessibility and POU5F1/SOX2 occupancy at these LTRs, whereas POU5F1 depletion reduced both SMARCA4 occupancy and chromatin accessibility, suggesting cooperative binding. Taxonomic classification showed that most SMARCA4-bound LTRs belong to evolutionary young, rodent-specific families. Human pluripotent cells were also analyzed for the binding of POU5F1, SOX2, and SMARCA4, which revealed that a similar POU5F1–SOX2–SMARCA4 architecture engages primate-specific LTR families distinct from mouse LTRs.

**Conclusions:** Our findings suggest that lineage-specific LTR repertoires can participate in a shared architecture of pluripotency TFs and chromatin remodeling factor. The association of evolutionarily distinct LTR repertoires with POU5F1/SOX2 and SMARCA4 in mouse and human pluripotent stem cells raises the possibility that non-orthologous, lineage-specific transposable elements were incorporated independently into similar regulatory architectures during mammalian evolution generating lineage-specific cis-regulatory networks.

## Background

Retrotransposons constitute approximately half of the mammalian genome (1, 2) and are now recognized as being integrated into host regulatory networks (3, 4). Their regulation has been characterized in pluripotent stem cells (PSCs), revealing that approximately half of the open chromatin regions overlap with retrotransposons (5, 6). Retrotransposons provide binding sites for pluripotency-related transcription factors (TFs), including POU5F1 (also called OCT4) and SOX2 (7–14). Mouse embryonic stem cells (mESCs) and rodent-specific long terminal repeat (LTR) retrotransposons, including RLTR9 and RLTR13D6, exhibit active regulatory features and are occupied by POU5F1 and SOX2 (11, 12). In human PSCs, primate- and human-specific LTR7 and LTR5_Hs elements exhibit active regulatory features and are targeted by core pluripotency-related TFs (10, 13, 14).

BAF/SWF-SNF chromatin remodeling complexes that contain SMARCA4 (also called BRG1) regulate chromatin accessibility (15, 16). In mESCs, SMARCA4 colocalizes with core pluripotency-related TFs including POU5F1 and SOX2 (17–21). Notably, POU5F1 facilitates the recruitment of SMARCA4 to its target sites, where SMARCA4 supports chromatin accessibility and POU5F1 binding (17). A recent study demonstrated that SMARCA4 binds to transposable elements (TEs) and contributes to TE-associated three-dimensional (3D) genome organization in human PSCs (22). However, whether SMARCA4 is preferentially recruited to specific TE families, how its occupancy is coupled to core pluripotency-related TFs, and whether such regulatory relationships are independently established in lineage-specific TE repertoires across mammalian species remain unclear.

In this study, we investigated how LTR retrotransposons engage the POU5F1–SOX2–SMARCA4 chromatin regulatory machinery in mESCs. We found that SMARCA4 selectively occupied evolutionarily recent *Mus*-specific LTR subfamilies that were co-occupied by POU5F1 and SOX2. SMARCA4 depletion was associated with reduced chromatin accessibility and POU5F1 and SOX2 occupancies at these LTRs, whereas POU5F1 depletion was accompanied by reduced SMARCA4 occupancy and chromatin accessibility. Moreover, analysis of human PSCs revealed that a similar POU5F1–SOX2–SMARCA4-associated chromatin architecture engages distinct LTR repertoires that are Primate-, Haplorrhini-, or Catarrhini-specific. Together, these findings suggest that during mammalian evolution, lineage-specific LTR repertoires were incorporated into the shared TF-chromatin remodeling architecture that shapes the 3D chromatin configuration in PSCs.

## Methods

### Datasets

All sequencing data analyzed in this study were obtained from the Gene Expression Omnibus (GEO) database. For mESCs, ChIP-seq data for BRG1 (SMARCA4), POU5F1 (OCT4), and SOX2 were retrieved from GSE87820, and ATAC-seq data were obtained from GSE87819 and GSE112130 datasets. ChIP-seq data for H3K27ac and H3K4me1 were obtained from GSE162739 and for H3K4me3 from GSE154379. For human PSCs, ChIP-seq data for POU5F1 and SOX2 were obtained from the GSE233635 dataset. ChIP-seq data for SMARCA4 and H3K27ac from human iPSCs were derived from GSE124903. H3K4me3 ChIP-seq data were from GSE201295. ATAC-seq datasets were obtained from GSE168002. RNA-seq data were from GSE87821. The details of accession numbers of the individual samples and their experimental group are summarized in Supplementary Table S1. The original publications describing these datasets are present in the reference list below (14, 17, 21, 23–26).

### ChIP-seq and ATAC-seq data analysis

Trim Galore (https://www.bioinformatics.babraham.ac.uk/projects/trim_galore) was used to remove adaptors and low-quality sequences. Clean reads were mapped to reference sequences (mm10 or hg38) using Hisat2 (27) with the options: --seed 100, --no-spliced-alignment, and --no-discordant. To ensure robust downstream analyses, high-quality reads mapped to a single genomic location were retained by filtering for an NH:i:1 tag and a mapping quality (MAPQ) score ≥60 using SAMtools (28). PCR duplicates were removed using SAMtools. Bigwig files were generated using deepTools BamCoverage (29) with the parameters --normalized using RPKM, --binsize 10. Peaks were identified along the reference sequences using MACS2 (30) with -f BAMPE option. The positions of the summits of the identified peaks were used in the subsequent analyses. The Deeptools compute matrix was used to create heat maps and aggregation plots. Plotting and statistical analyses were performed using R 4.3.1 software unless otherwise indicated.

### Transposable element enrichment analysis

Enrichment analysis was performed locally using Transposable Element Enrichment Analyzer (TEENA) (31) with default parameters and promoter regions retained. The SMARCA4 peak summits were used. Human and mouse datasets were analyzed using the hg38 and mm10 genome assemblies, respectively. For human analyses, the hg38 reference genome sequence, chromosome size, and RepeatMasker annotations were obtained from the UCSC Genome Browser and gene annotations were obtained from Ensembl Release 110 (Homo_sapiens.GRCh38.110.gtf). For mouse analyses, the mm10 reference genome sequence, chromosome size, and RepeatMasker annotations were obtained from the UCSC Genome Browser, and gene annotations corresponding to GRCm38 were obtained from Ensembl Release 102 (Mus_musculus.GRCm38.102.gtf). Enrichment of individual TE subfamilies was evaluated using Fisher’s exact test implemented in TEENA, followed by Benjamini–Hochberg correction for multiple testing. Fold-enrichment values and adjusted p values generated by TEENA were used for downstream analysis and visualization.

### Structural classification of TEs

The genomic structures of the SMARCA4-bound TEs were classified by manual inspection of the RepeatMasker annotations surrounding each locus. LTR elements lacking an associated endogenous retrovirus (ERV) internal region were classified as solo LTRs, whereas ERV internal sequences lacking a recognizable pair of corresponding flanking LTRs were classified as internal fragments. Loci located within a recognizable LTR–internal–LTR structure, based on the compatibility of the LTR and internal ERV annotations, orientation, and genomic organization, were classified as the proviral form. Nested insertion of unrelated TEs was permitted when the underlying proviral organization remained unambiguous. Loci with complex or ambiguous arrangements that could not be confidently assigned to these categories were classified as undetermined or complex.

### Transposable element expression analysis

The method to calculate transposon expression levels have been described in a previous study (9). Briefly, mRNA sequence reads were mapped to the mouse reference genome mm10 using STAR (32) with the options --outFilterMultimapNmax 100 and --winAnchorMultimapNmax 200 to retain multimapped reads. Subsequently, TEcount (33) was used with multimode to quantitate expression levels of TE subfamilies. Differential expression analysis was conducted using DESeq2 (34) on TEcount-generated raw count reads containing both gene- and TE-mapped reads. Differential expressions were identified using thresholds of adjusted p < 0.05 and difference was 1.5-fold or more. For bar-plot visualization, values were normalized to reads per million (RPM) with the denominator defined as the sum of reads mapping to genes and to TEs.

### Taxonomic classification of transposable elements

Taxonomic classifications of retrotransposon families were obtained from Dfam (35) using FamDB. RepeatMasker subfamily names were queried against the local Dfam FamDB database. The resulting taxonomic assignments were manually inspected to confirm consistency with the Dfam annotations. Taxonomic assignments were used to characterize the lineage distribution of the LTR subfamilies rather than to infer precise insertion ages.

### Motif analysis

For motif enrichment analysis, Analysis of Motif Enrichment (AME) tool (36) was used to assess motif enrichment under default parameters. JASPAR CORE (2026) was selected in “Input the motifs.” The presence of POU5F1::SOX2 motifs within the sequences were examined using Find Individual Motif Occurrences (FIMO) (37). The Motif matrix MA0142.1, which represented POU5F1::SOX2, was downloaded from the JASPAR database (https://jaspar.genereg.net).

### Sequence divergence analysis

Sequence divergence values of individual LTR copies were obtained from the NCBI Genome Browser (mm10.rmsk.txt). Sequence divergence from the corresponding RepeatMasker consensus sequence was evaluated using the MilliDiv value in the RepeatMasker annotation and converted to percent divergence.

### A/B compartment analysis

The A/B chromatin compartment information for mESCs was obtained from a publicly available Hi-C dataset (GSE164068) (38). The processed PC1 bedGraph file (GSE164068_ESC_LMNA_ctr_merge_mm10.PC1.bedGraph.gz) was downloaded from the GEO database. Following the original study, regions with positive PC1 values were classified as A compartments, whereas regions with negative PC1 values were classified as B compartments.

## Result

### SMARCA4 selectively occupied specific LTR subfamilies in mESCs

To investigate whether SMARCA4 occupancy was associated with specific retrotransposons in mESCs, we analyzed the published SMARCA4 ChIP-seq data (17). Only high-quality reads that uniquely mapped to the genome were used for peak calling (see Methods section). Genome-wide overlap analysis using TEENA (31) revealed that of 5,181 SMARCA4 ChIP-seq peak summits, 994 (19%) were located inside retrotransposons. Among these, 43 LTR retrotransposon subfamilies, including RLTR13D6, RLTR9E, and RLTR9D, were enriched significantly (adjusted p < 0.05) for SMARCA4 occupancy (Fig. 1A). The top 10 enriched TE subfamilies are shown in Fig. 1A, and a complete list of the enriched TE subfamilies is provided in Supplementary Table S2. Most of the enriched LTR elements (40 subfamilies) were LTR sequences, whereas a minority (three subfamilies) were internal regions. Seven SINE B1 subfamilies were enriched, whereas none of the B2 subfamilies was enriched. We did not detect the enrichment of LINE subfamilies. Therefore, SMARCA4 preferentially binds to LTR sequences of LTR retrotransposons. RLTR9E is an LTR sequence of MMERVK9E that is distantly related to the RLTR13 subfamilies in the tree of ERVK class LTRs (6). However, both RLTR13 subfamilies and RLTR9-related LTRs (RLTRETN_Mm, ERVB4_1C-LTR, and ERVB7_1-LTR_MM) constitute the LTR sequences of MusD (ETnERV-int) and ETn (MMETn-int) (6), the latter of which is transcribed in mESCs (39). Structural classification (see Methods section) revealed that 677 (86.1%) of the 786 SMARCA4-associated LTR retrotransposon loci were the solo LTRs. In contrast, 22 loci (2.8%) showed a proviral structure, 16 (2.0%) were internal fragments, and 71 (9.0%) were classified as undetermined or complex (Fig. S1).

**Fig1.**
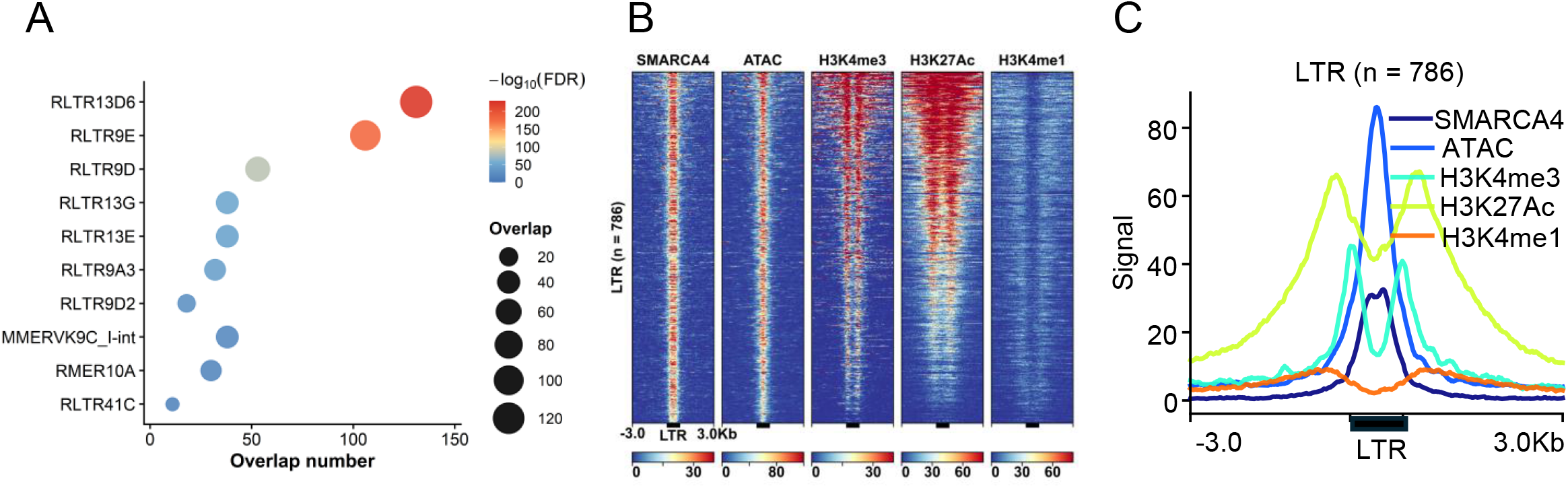
SMARCA4 preferentially occupies specific LTR retrotransposon subfamilies in mouse embryonic stem cells. (A) Enrichment of TE subfamilies of SMARCA4 binding in mouse embryonic stem cells. The top 10 significantly enriched TE subfamilies are shown. Dot size indicates the number of overlapping SMARCA4 peaks, and color indicates the −log_10_(FDR). **(B and C)** Mapped reads (normalized as reads per kilobase per million mapped reads; RPKM) from SMARCA4 ChIP-seq, ATAC-seq, H3K4me3, H3K27ac, and H3K4me1 ChIP-seq data are shown as tornado plots (B) and aggregated line plots (C). LTRs in the heatmaps are ordered by SMARCA4 signal intensity.

Next, we examined the chromatin states of SMARCA4-enriched LTRs. Analysis of published ChIP-seq and ATAC-seq datasets in mESCs (17, 24, 40) revealed that SMARCA4-bound LTRs exhibit chromatin accessibility and H3K4me3 and H3K27Ac signals, although they show little H3K4me1 signal (Fig. 1B and 1C). Among the LTR subfamilies, RLTR13D6 and RLTR9E showed particularly strong enrichment of SMARCA4 and were associated with active chromatin states in mESCs. Thus, our results are consistent with those of previous studies (11, 12, 41, 42) and reveal extensive SMARCA4 occupancy at these LTR subfamilies.

### POU5F1 and SOX2 bound to SMARCA4-associated LTR subfamilies

To elucidate the sequence characteristics of SMARCA4-bound LTRs, we performed motif enrichment analysis using AME (36), which identified enrichment of the POU5F1(OCT4)-SOX2 binding motif (Fig. 2A). RLTR13D6 and RLTR9E, two LTR subfamilies showing strong SMARCA4 enrichment, contained POU5F1::SOX2 motif matches in their consensus sequences (Fig. S2). Analysis of published POU5F1 and SOX2 ChIP-seq data (17) showed that POU5F1 and SOX2 bind to SMARCA4-occupied LTRs (Fig. 2B). The ChIP-seq profiles resembled those of SMARCA4 ChIP-seq, suggesting co-binding (Fig. 2C). The co-localization of POU5F1 and SOX2 with accessible, H3K27ac-enriched chromatin at RLTR9E and RLTR13D6 is consistent with previous studies identifying these LTR subfamilies as active regulatory elements in mESCs (11, 12). Importantly, our analysis revealed that SMARCA4 is enriched at these POU5F1/SOX2-occupied LTRs. Together, these findings suggest that specific LTR subfamilies provide genomic platforms where core pluripotency TFs and SMARCA4 chromatin-remodeling machinery converge.

**Fig.2.**
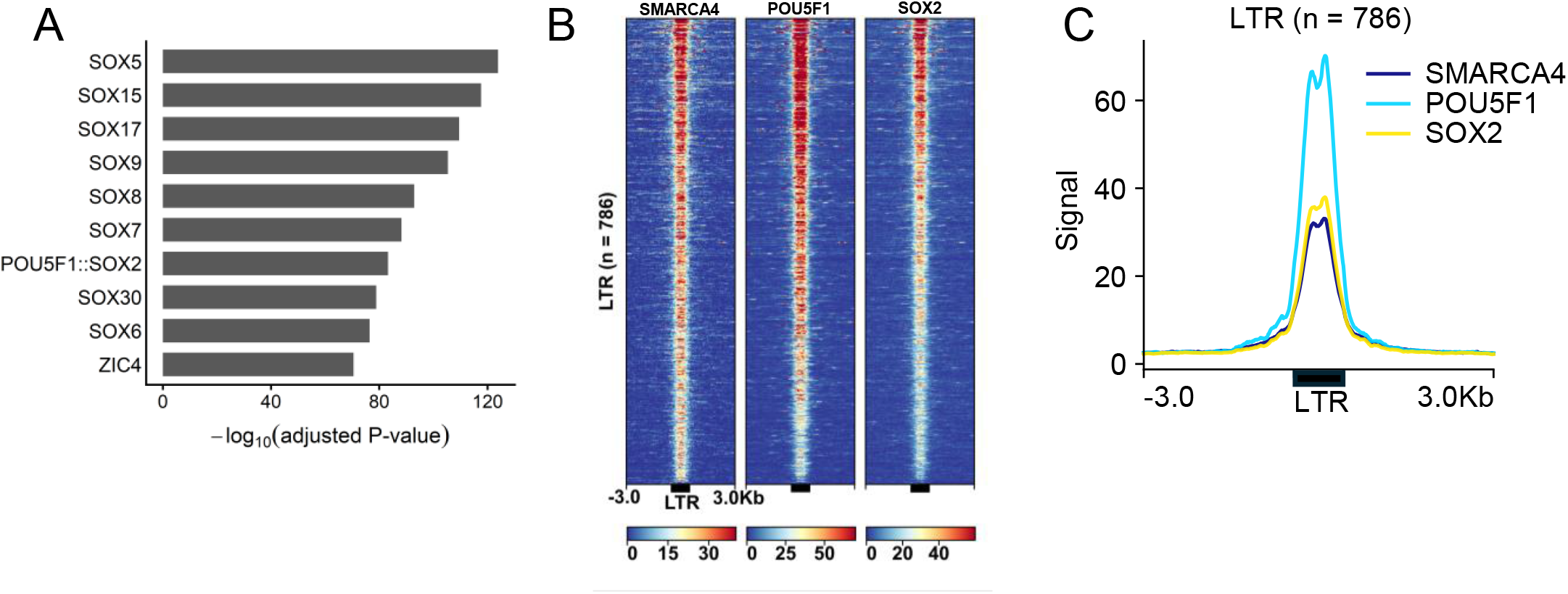
POU5F1 and SOX2 bind at SMARCA4-bound LTRs. (A) Enrichment of transcription factor-binding motifs in SMARCA4-bound LTR sequences. The top 10 significantly enriched motifs are shown. **(B and C)** Mapped reads (normalized as reads per kilobase per million mapped reads; RPKM) from SMARCA4, POU5F1, and SOX2 ChIP-seq data are shown as tornado plots **(B)** and aggregated line plots **(C)**. LTRs in the heatmaps are ordered by SMARCA4 signal intensity.

### SMARCA4 supports chromatin accessibility and POU5F1/SOX2 occupancy at LTR elements

SMARCA4 plays an important role in maintaining chromatin accessibility in PSCs (17). Therefore, we next explored whether SMARCA4 association with these LTRs is functionally required for an accessible chromatin state. To investigate the effect of SMARCA4 loss, we reanalyzed published ATAC-seq data from SMARCA4 conditional-knockout mESCs (17). In this system, SMARCA4 deletion was induced by tamoxifen (TAM)-mediated activation of CreER. We compared untreated and TAM-treated cells to assess the changes associated with SMARCA4 depletion. SMARCA4 depletion resulted in reduced ATAC-seq signals in SMARCA4-bound LTRs (Fig. 3A–C). To further validate this reduction in chromatin accessibility of LTRs, we analyzed an independent publicly available ATAC-seq dataset under SMARCA4 wild-type and knockout conditions (23). Again, the chromatin accessibility of SMARCA4-bound LTRs was downregulated (Fig. 3D–F).

**Fig.3.**
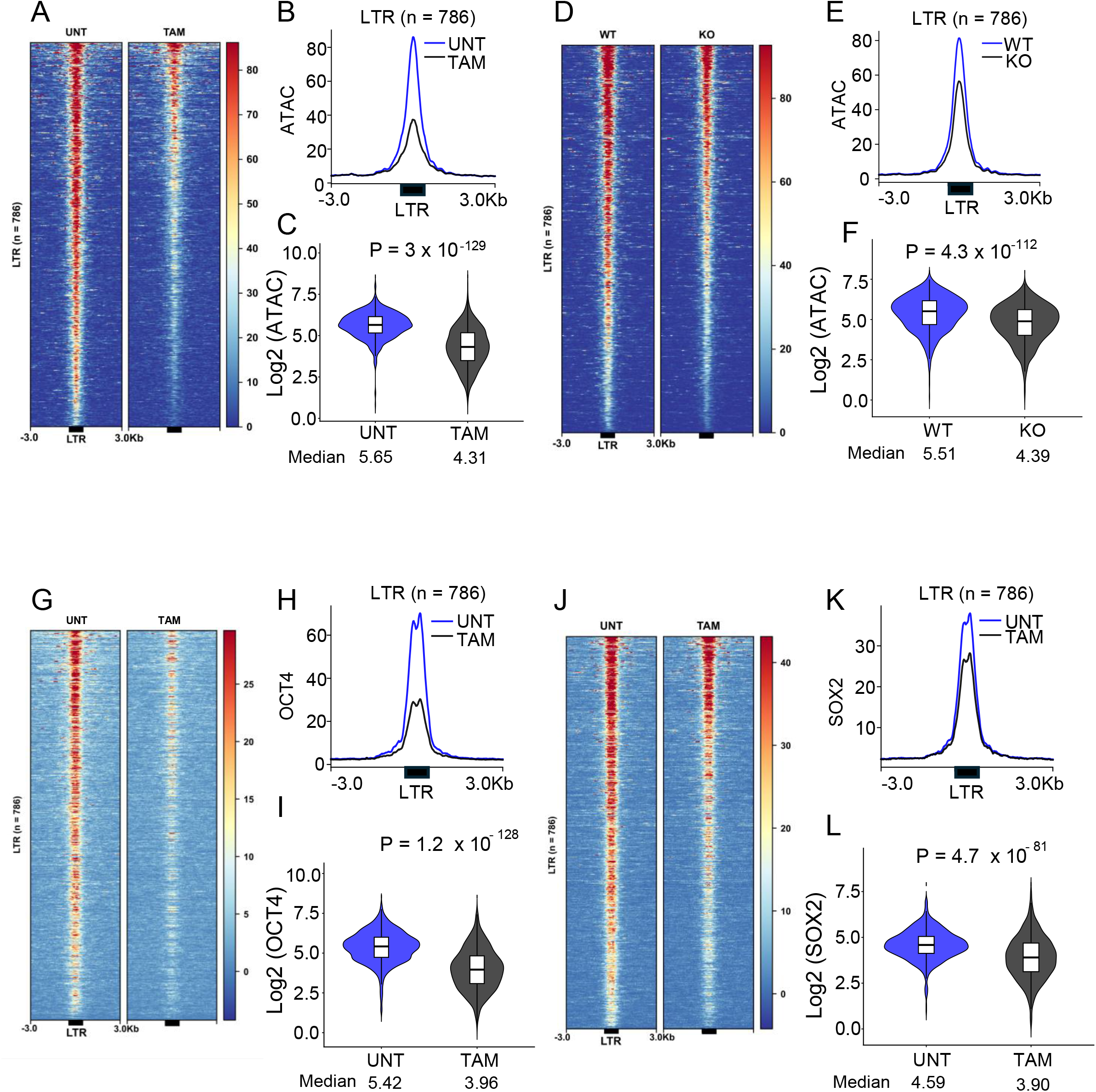
SMARCA4 depletion reduces chromatin accessibility and POU5F1/SOX2 occupancy at SMARCA4-bound LTRs. **(A–C)** ATAC-seq signals at SMARCA4-bound LTRs (n = 786) in untreated (UNT) and tamoxifen-treated (TAM) mESCs. **(D–F)** ATAC-seq signals at SMARCA4-bound LTRs in SMARCA4 wild-type (WT) and knockout (KO) mESCs. **(G–I)** POU5F1 ChIP-seq signals and **(J–L)** SOX2 ChIP-seq signals at SMARCA4-bound LTRs in UNT and TAM-treated mESCs. Mapped reads (normalized as reads per kilobase per million mapped reads; RPKM) are shown as tornado plots **(A, D, G, and J)**, aggregated line plots **(B, E, H, and K)**, and violin plots **(C, F, I, and L)**. LTRs in all tornado plots are ordered by SMARCA4 signal intensity. *P* values in the violin plots were calculated using the Wilcoxon rank-sum test.

Next, we analyzed POU5F1 and SOX2 ChIP-seq datasets under SMARCA4-depleted conditions (17) to determine whether SMARCA4 loss also affected TF binding at these LTRs. POU5F1 and SOX2 occupancy was reduced following SMARCA4 depletion at the target LTRs (Fig. 3G–L). Since the reduction of SMARCA4 did not result in a loss of pluripotency or reduced expression of POU5F1 and SOX2 (17), these results suggest that SMARCA4 is important for the binding of POU5F1 and SOX2 to LTR sequences and for creating open chromatin.

### POU5F1 is required for SMARCA4 occupancy and chromatin accessibility at target LTRs

Previous studies have demonstrated that POU5F1 can facilitate SMARCA4 recruitment at its binding sites, suggesting a relationship between pluripotency-related TFs and chromatin remodeling (17). Therefore, we investigated whether POU5F1 is required for SMARCA4 occupancy of LTRs. To address this, we reanalyzed the ATAC-seq and SMARCA4 ChIP-seq datasets under POU5F1 depletion (17). This analysis revealed a reduction in both chromatin accessibility and SMARCA4 occupancy at the target LTRs (Fig. 4), suggesting that POU5F1 contributes to the maintenance of chromatin accessibility and SMARCA4 occupancy at the target LTRs, whereas SMARCA4 supports both accessibility and POU5F1/SOX2 occupancy. Taken together, these findings suggest a mutually reinforcing relationship in which POU5F1 and SMARCA4 contribute to the maintenance of active chromatin states at specific LTR elements.

**Fig.4.**
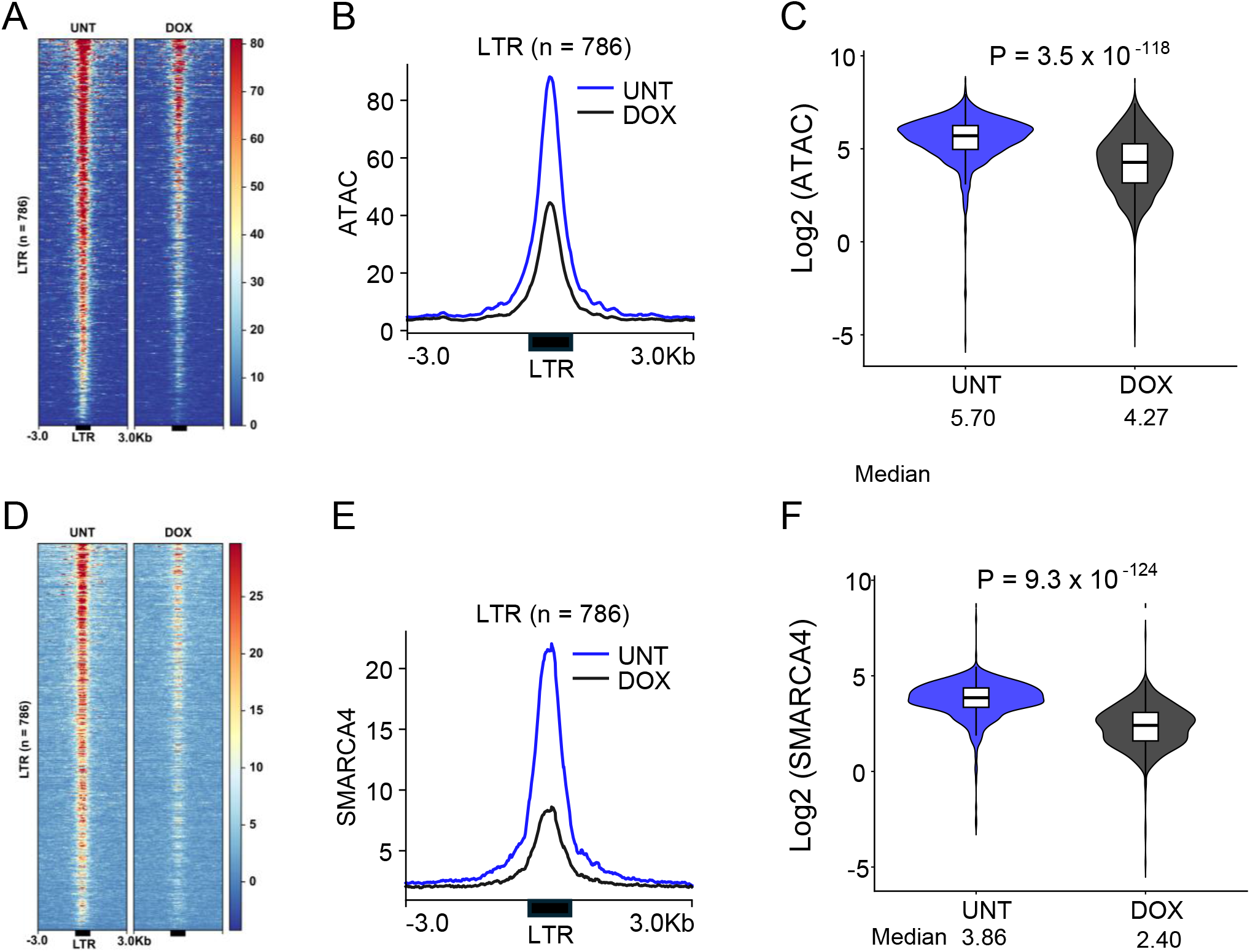
POU5F1 depletion reduces chromatin accessibility and SMARCA4 occupancy at SMARCA4-bound LTRs. **(A–C)** ATAC-seq signals at SMARCA4-bound LTRs (n = 786) in untreated (UNT) and doxycycline-treated (DOX) mESCs. **(D–F)** SMARCA4 ChIP-seq signals at SMARCA4-bound LTRs in UNT and DOX-treated mESCs. Mapped reads (normalized as reads per kilobase per million mapped reads; RPKM) are shown as heatmaps **(A and D)**, aggregated line plots **(B and E)**, and violin plots **(C and F)**. LTRs in all heatmaps are ordered by SMARCA4 signal intensity. *P* values in the violin plots were calculated using the Wilcoxon rank-sum test.

### SMARCA4 and POU5F1 depletion differentially affected TE expression in mESCs

Next, we examined whether SMARCA4 and POU5F1 depletion affected TE expression using publicly available RNA-seq datasets (17). SMARCA4 depletion resulted in a few differentially expressed TE subfamilies, whereas POU5F1 depletion led to significant downregulation of 21 TE subfamilies (Fig. S3). Among the downregulated TEs were MMETn-int and ETnERV-int, which are highly expressed in mESCs (39). Notably, RLTR13G, an LTR subfamily associated with the ERV family (1), was downregulated following POU5F1 depletion. RLTR13G was enriched in SMARCA4 occupancy and bound POU5F1 (Fig. 1 and 2), which was consistent with the observed reduction in its expression upon POU5F1 depletion. These findings suggested that POU5F1 contributes to the transcriptional activity of specific TE subfamilies, including those associated with SMARCA4-bound LTRs, whereas SMARCA4 depletion has a relatively limited effect on TE expression.

### SMARCA4-associated LTRs are enriched for LTR repertoires associated with relatively recent evolutionary lineages

As retrotransposons have expanded in different eras during mammalian evolution (43, 44), we next examined whether SMARCA4 binding to the selected LTRs reflects their evolutionary ages. To explore this, we assigned individual LTR subfamilies to their respective taxonomic lineages based on Dfam annotations (35) and compared the composition of SMARCA4-bound LTRs with that of the genomic LTR repertoire. Compared to the overall LTRs in the mouse genome, SMARCA4-bound LTRs were biased toward relatively young Rodentia-specific subfamilies (Fig. 5A and 5B). Next, we examined whether the lineage bias was driven by a single dominant LTR subfamily. Although RLTR13D6 and RLTR9E provided relatively large portions of SMARCA4-bound copies, SMARCA4 occupancy was distributed across multiple rodent-specific LTR subfamilies, including RLTR9D, RLTR13G, and RLTR13E (Fig. 5C). Thus, lineage bias among SMARCA4-bound LTRs does not reflect the contribution of a single LTR subfamily, but rather the contributions of multiple LTR subfamilies associated with related evolutionary lineages. Together, these results reveal that the POU5F1–SMARCA4 regulatory architecture in mESCs preferentially engages a lineage-specific LTR repertoire, raising the possibility that distinct LTR families may have been recruited into this regulatory network in mouse/rodent-associated lineages.

**Fig.5.**
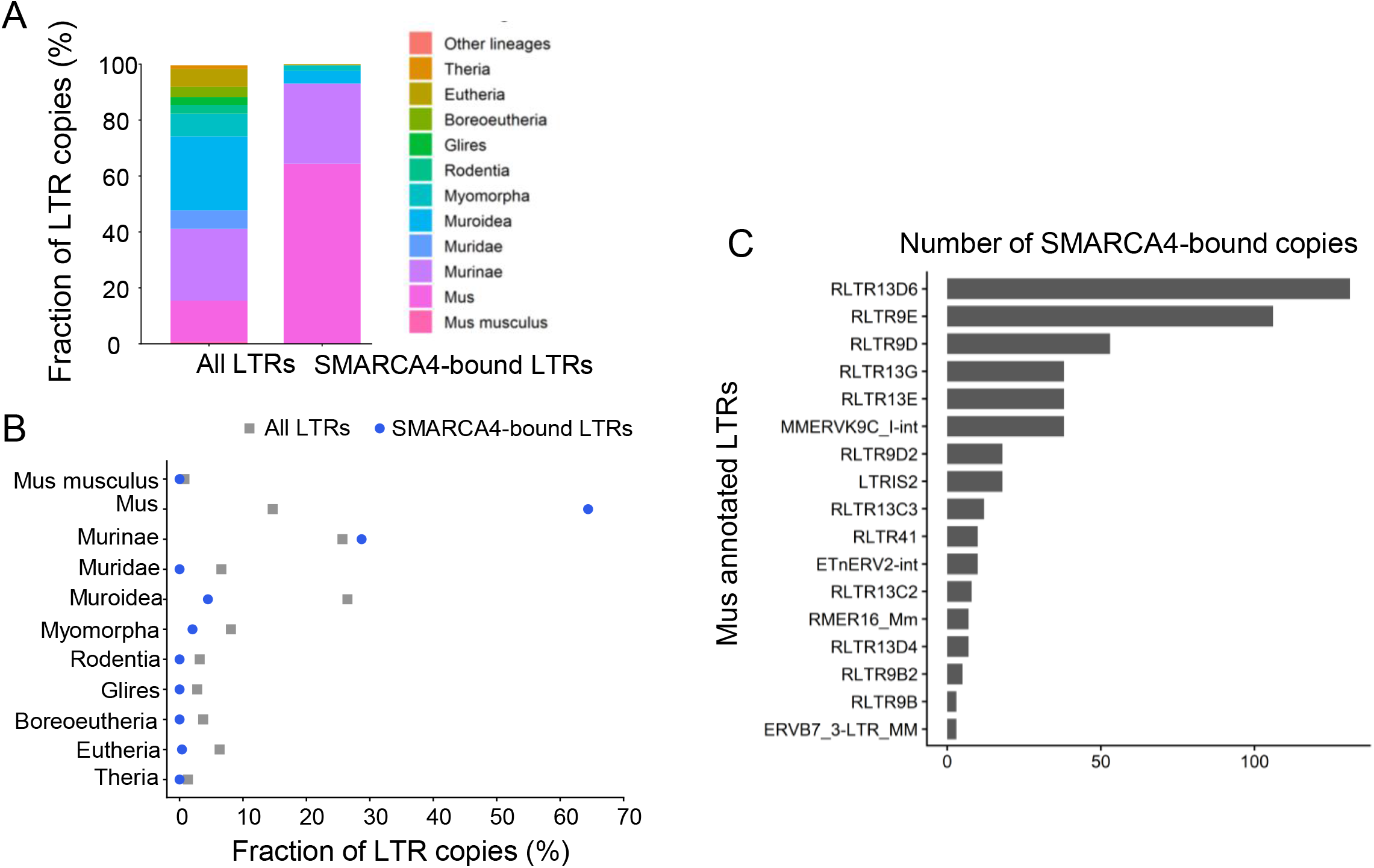
SMARCA4-bound LTRs are biased toward LTR repertoires associated with relatively recent evolutionary lineages. **(A)** Taxonomic lineage distribution of all annotated LTRs and SMARCA4-bound LTRs in the mouse genome. LTR subfamilies were assigned to taxonomic lineages based on Dfam annotations. **(B)** Fractions of all annotated LTRs and SMARCA4-bound LTRs assigned to each taxonomic lineage. **(C)** Number of SMARCA4-bound copies for individual LTR subfamilies.

We further explored the characteristics of SMARCA4-bound and unbound copies within the same LTR subfamily. SMARCA4-bound copies showed higher chromatin accessibility, active histone modification signals, and POU5F1/SOX2 occupancy than the unbound copies (Fig. S4A and S4B). However, the distribution of sequence divergence between the SMARCA4-bound and unbound copies of both RLTR13D6 and RLTR9E was similar (Fig. S4C and S4D). POU5F1::SOX2 motifs were detected in nearly all the copies, irrespective of SMARCA4 occupancy (Fig. S4E and S4F). We also examined the A/B compartment distribution of SMARCA4-bound and unbound copies. RLTR13D6 showed similar compartment distributions between SMARCA4-bound and unbound copies, whereas SMARCA4-bound RLTR9E copies showed a higher proportion in compartment A than unbound copies (Fig. S4G). These results suggest that SMARCA4 occupancy at individual RLTR13D6 and RLTR9E copies cannot be explained by sequence divergence or the presence of the POU5F1::SOX2 motif, whereas its association with the A/B compartment distribution appears to differ between LTR subfamilies.

### Distinct LTR subfamilies engage a similar POU5F1–SMARCA4-associated chromatin architecture in human pluripotent cells

Finally, we asked whether a similar regulatory architecture is constructed in LTRs in human PSCs and, if so, whether it engages a distinct set of LTRs because RLTR13, RLTR9, and other SMARCA4-bound LTRs are rodent-specific and thus absent in the human genome. To address this, we analyzed the published SMARCA4 ChIP-seq data from human iPSCs (26). As observed in the mESCs, SMARCA4 occupancy was enriched in specific LTR subfamily members (Fig. 6A). A complete list of the enriched TE subfamilies is provided in Supplementary Table S2. Motif enrichment analysis identified enrichment of the POU5F1::SOX2 motif in SMARCA4-bound LTRs (Fig. 6B), as observed for mouse SMARCA4-bound LTRs. LTR49 and MER4D, two LTR subfamilies that showed strong SMARCA4 enrichment, contained POU5F1::SOX2 motif matches in their consensus sequences (Fig. S5). Analysis of published ChIP-seq datasets of human PSCs (14, 25, 26) showed that SMARCA4-bound human LTRs exhibited chromatin accessibility and active histone modifications, including H3K4me3 and H3K27ac, and were co-occupied by POU5F1 and SOX2 (Fig. 6C and 6D), as observed for SMARCA4-bound mouse LTRs in mESCs.

**Fig.6.**
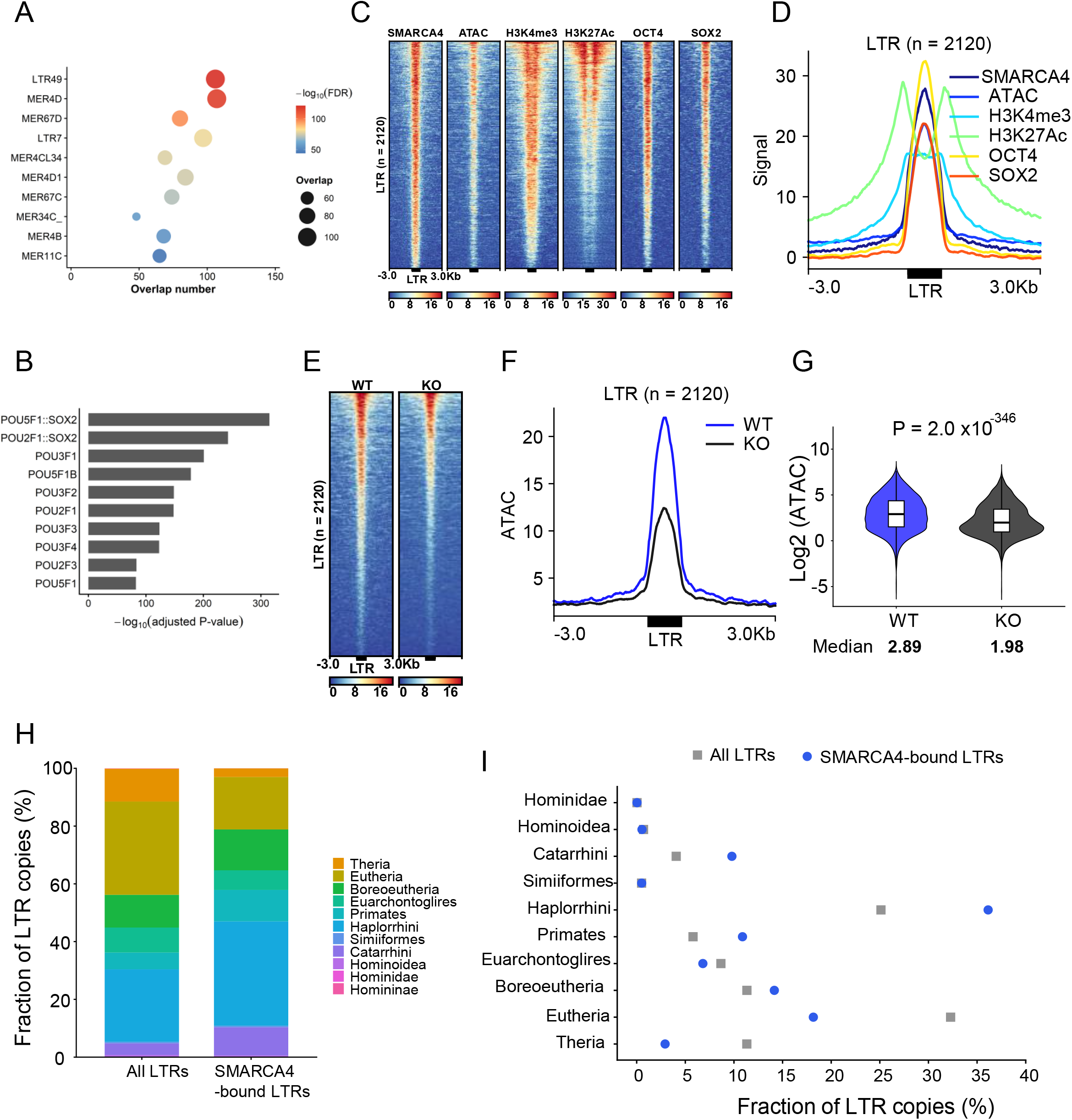
SMARCA4-associated LTRs engage POU5F1/SOX2-associated chromatin architecture in human PSCs. **(A)** Enrichment of TE subfamilies at SMARCA4-binding sites in human PSCs. The top 10 significantly enriched TE subfamilies are shown. Dot size indicates the number of overlapping SMARCA4 peaks, and color indicates the −log10(FDR). **(B)** Enrichment of transcription factor-binding motifs in SMARCA4-bound LTR sequences. The top 10 significantly enriched motifs are shown. **(C and D)** Mapped reads (normalized as reads per kilobase per million mapped reads; RPKM) from SMARCA4, ATAC-seq, H3K4me3, H3K27ac, POU5F1, and SOX2 datasets are shown as heatmaps **(C)** and aggregated line plots **(D)** at SMARCA4-bound LTRs (n 2120). LTRs in the heatmaps are ordered by SMARCA4 signal intensity. **(E–G)** ATAC-seq signals at SMARCA4-bound LTRs in SMARCA4 wild-type (WT) and knockout (KO) cells, shown as a heatmap **(E)**, aggregated line plot **(F)**, and violin plot **(G)**. The *P* value in **(G)** was calculated using the Wilcoxon rank-sum test. **(H)** Taxonomic lineage distribution of all annotated LTRs and SMARCA4-bound LTRs in the human genome. **(I)** Fractions of all annotated LTRs and SMARCA4-bound LTRs assigned to each taxonomic lineage.

Next, we examined whether chromatin accessibility at these LTRs was dependent on SMARCA4, similar to that in the mouse model. Analysis of ATAC-seq data from wild-type and SMARCA4 knockout human PSCs revealed reduced chromatin accessibility following SMARCA4 depletion (Fig. 6E–G). These findings indicate that the SMARCA4-dependent maintenance of accessible chromatin at LTRs is shared between mouse and human PSCs. Although POU5F1 occupancy following SMARCA4 depletion could not be assessed because the corresponding datasets were unavailable, these results suggest a similar SMARCA4-associated chromatin architecture at LTRs in human and mouse PSCs.

Examination of the taxonomic lineage distribution of SMARCA4-bound LTRs in human PSCs revealed that the LTR subfamilies of primate-related evolutionary lineages (Catarrhini-, Haplorrhini-, and primate-specific) were enriched (Fig. 6H and 6I). These findings suggest that distinct lineage-specific LTR repertoires engage in a shared pluripotency-related TF–chromatin remodeling architecture in mouse and human PSCs.

## Discussion

It has been shown that, while the sequence motif in the genomic DNA is essential for the binding of pioneer TFs such as POU5F1 to bind to the DNA, the chromatin remodeling factor SMARCA4 is required for recruiting other TFs and consequent gene activation (17). SMARCA4 is a SWI/SNF-type ATPase of the BAF chromatin remodeling complex that can remodel nucleosome positions and even transfer nucleosomes between DNAs (15, 45). Therefore, the collaborative actions of pioneer TFs and chromatin remodelers determine open chromatin regions and robust cell-type-specific transcriptional states. This study showed that a substantial fraction of POU5F1–SOX2–SMARCA4-mediated open chromatin regions in mouse and human PSCs are provided by selective LTR retrotransposons, such as the mouse RLTR13 and RLTR9 subfamilies and human LTR49 and MER4D. Recently, it was reported that SMARCA4 promotes TE-associated 3D genome organization in human PSCs, although the specific TE has not yet been identified (22). Therefore, it is conceivable that POU5F1–SOX2–SMARCA4-bound LTR retrotransposons contribute to the regulation of 3D genome organization, which likely underlies the transcriptomic states in early embryogenesis in humans and mice.

Notably, these LTR subfamilies are evolutionary young and specific to the mouse and human lineages, respectively. Therefore, their insertion sites in the mouse and human genomes are not orthologous, and their downstream effects on gene expression are likely different. Thus, conserved host regulatory mechanisms, including POU5F1/SOX2 and SMARCA4, seem to have been repeatedly engaged in newly available TE-derived sequences in each lineage during mammalian evolution, generating diverse regulatory networks. In totipotent cells, mouse MT2/MERVL and human HERVL/MLT2 serve as binding sites for pioneer TFs to play regulatory roles in zygotic genome activation, thereby establishing transcriptomes for the initiation of embryogenesis (46, 47). MERVL and HERVL were inserted into non-orthologous loci in mouse and human genomes, respectively. In contrast, the binding sites of pioneer TFs in human neural stem cells and breast cancer-derived cells are mainly provided by ancient retrotransposon families (43, 48), suggesting that orthologous loci have conserved gene regulatory functions across mammalian species during the developmental stages of organogenesis. These results led us to the idea that the diversity of retrotransposon sequences and their insertion sites in different taxonomic clades underlies the *hourglass model* of ontogenesis (49), where morphology in early embryos is diversified among animals, but conserved in organogenesis stages, and again becomes diversified in the late stage of development to produce species-specific morphological traits. The diversity of the transcriptomic trajectories during development follows this model (50). Therefore, it is tempting to propose that retrotransposon transpositions underlie transcriptomic diversity in early embryogenesis.

From a view of TEs, carrying binding sites of pluripotency-related TFs is likely advantageous for their persistence in the host genome. As random mutations accumulate within TE sequences over time and eventually inactivate TEs, active transposition is essential for TE survival. To be inherited by subsequent generations, such TE transposition must take place in germ cells or in developmental stages before germ-cell specification. We showed that LTRs of ETn and MusD carry POU5F1/SOX2/SMARCA4 binding sites to drive transcription, which is consistent with their persistence and active transposition in the evolutionarily recent mouse genome (51, 52).

LTR retrotransposons are present in two forms. Upon retrotransposition, the new copy is in a full-length form with 5’ LTR, the internal sequence, and 3’ LTR. These full-length copies occasionally undergo homologous recombination between the two LTRs, resulting in a single LTR sequence called solo LTR. We found that most LTR loci that bound to SMARCA4 were solo LTRs, suggesting relaxed epigenetic silencing of solo LTRs. This is similar to the case observed in male mouse germ cells, where DNA-hypomethylated copies of intracisternal A-particle (IAP) retrotransposons are preferentially solo LTRs (53, 54).

Motif presence alone did not appear to determine SMARCA4 occupancy in the individual LTR copies. In RLTR13D6 and RLTR9E, POU5F1::SOX2 motif matches were detected in many copies, whereas only a subset of copies were occupied by SMARCA4 and TFs. In addition, the sequence divergence and localization of the A/B compartment did not clearly separate the SMARCA4-bound and unbound copies. This interpretation is consistent with experimental studies on HERV-H-derived sequences in human PSCs. A previous study showed that *de novo* insertion of HERV-H elements could create new boundaries of topologically associating domains; however, boundary formation depended on their insertion sites (55). Thus, TE-derived sequences have regulatory or architectural potential; however, this potential is realized only in permissive genomic and chromatin contexts. Similarly, our results indicated that many LTR copies contained POU5F1/SOX2 motif potential, but only selected copies were incorporated into SMARCA4-associated active chromatin states.

Our study had some limitations. Dfam-based taxonomic assignment was used to describe the lineage distribution of LTR subfamilies; however, it did not provide precise insertion ages for individual copies. Thus, our evolutionary analysis should be considered at the level of TE subfamily distribution rather than the exact locus-level insertion ages. In addition, our analysis underestimated the function of young TE copies with highly similar sequences because our analyses relied on uniquely mapped reads. Therefore, the contribution of the recently expanded TE repertoires may have been conservatively estimated in this study. However, our findings are consistent with a model in which lineage-specific LTR retrotransposons provide genomic platforms for the shared POU5F1/SOX2–SMARCA4 chromatin regulatory architecture in PSCs. More broadly, our study highlighted how evolutionarily distinct TE repertoires can be incorporated into the conserved host regulatory machinery, thereby contributing to the establishment of cell type-specific transcriptomes in a species-specific manner.

## Conclusions

In this study, we showed that specific LTR retrotransposon subfamilies engage with the POU5F1/SOX2–SMARCA4 chromatin regulatory machinery in PSCs. In mESCs, SMARCA4 preferentially occupies the lineage-biased LTR repertoires associated with chromatin accessibility and POU5F1/SOX2 occupancy. Conversely, POU5F1 depletion reduced SMARCA4 occupancy and chromatin accessibility at these LTRs, supporting a mutually reinforcing relationship between pluripotency TFs and chromatin remodeling machinery. A similar regulatory architecture is observed in human pluripotent cells, but engages in distinct LTR repertoires. Together, these findings suggest that, during mammalian evolution, lineage-specific LTR repertoires can be incorporated into the shared architecture comprised by TFs and chromatin remodeling factors and can contribute to lineage-specific regulatory innovation in the early stages of embryonic development.

## Supporting information

All Supplemental Figures

All Supplemental Tables

## List of abbreviations

LTR: long-terminal repeat
mESC: mouse embryonic stem cell
PSC: pluripotent stem cell
TAD: topologically associated domain
TE: Transposable element
TF: transcription factor
ERV: endogenous retrovirus

## Declarations

The authors declare no conflict of interests.

## Funding

Ministry of Education, Culture, Sports, Science, and Technology (MEXT), Japan, to H. S. 2825KJ1422.

Ministry of Education, Culture, Sports, Science, and Technology (MEXT), Japan, to K. I. 25H01299, 23K18153, and 19K22452.

## Availability of data and materials

All sequencing data in this study are publicly available, and their accession numbers are indicated in the Methods section and Supplementary Tables. The codes used in the analyses can be obtained from the authors upon request.

## Acknowledgements

1. H. S. was supported by the Make New Standards Program at Nagoya University and JSPS DC Research Fellowships from MEXT, Japan. K. I. was supported by a Grant-in-Aid for Challenging Exploratory Research and a Grant-in-Aid for Transformative Research Areas from MEXT, Japan. We would like to thank Editage (www.editage.jp) for the English language editing.

## Author information

### Contributions

H. S. designed the study, analyzed the data, and wrote the manuscript. K. I. supervised the study and revised the manuscript.

### Corresponding authors

Kenji Ichiyanagi and Hikaru Suzuki.

### Ethics approval and consent to participate

Not applicable

### Consent for publication

All authors have reviewed the manuscript and consented to its publication.

