## Supplementary material for "Lineage-specific LTR Retrotransposon Repertoires Engage OCT4–SOX2–SMARCA4 Chromatin Regulatory Architecture in Mouse and Human Pluripotent Stem Cells": All Supplemental Figures

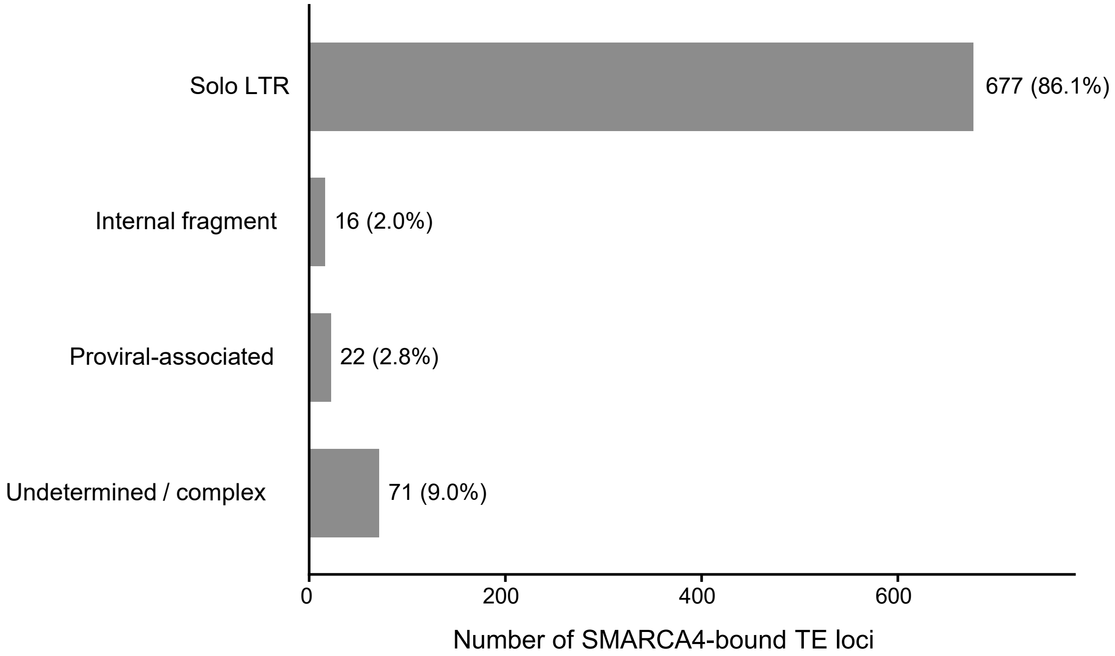


**Fig.S1 Structural classification of SMARCA4-bound TE loci.**

Structural classification of SMARCA4-bound TE loci (n = 786) based on RepeatMasker annotations. The number and percentage of loci classified as solo LTR, internal fragment, proviral-associated, or undetermined/complex are shown.

Suzuki_Fig. S2

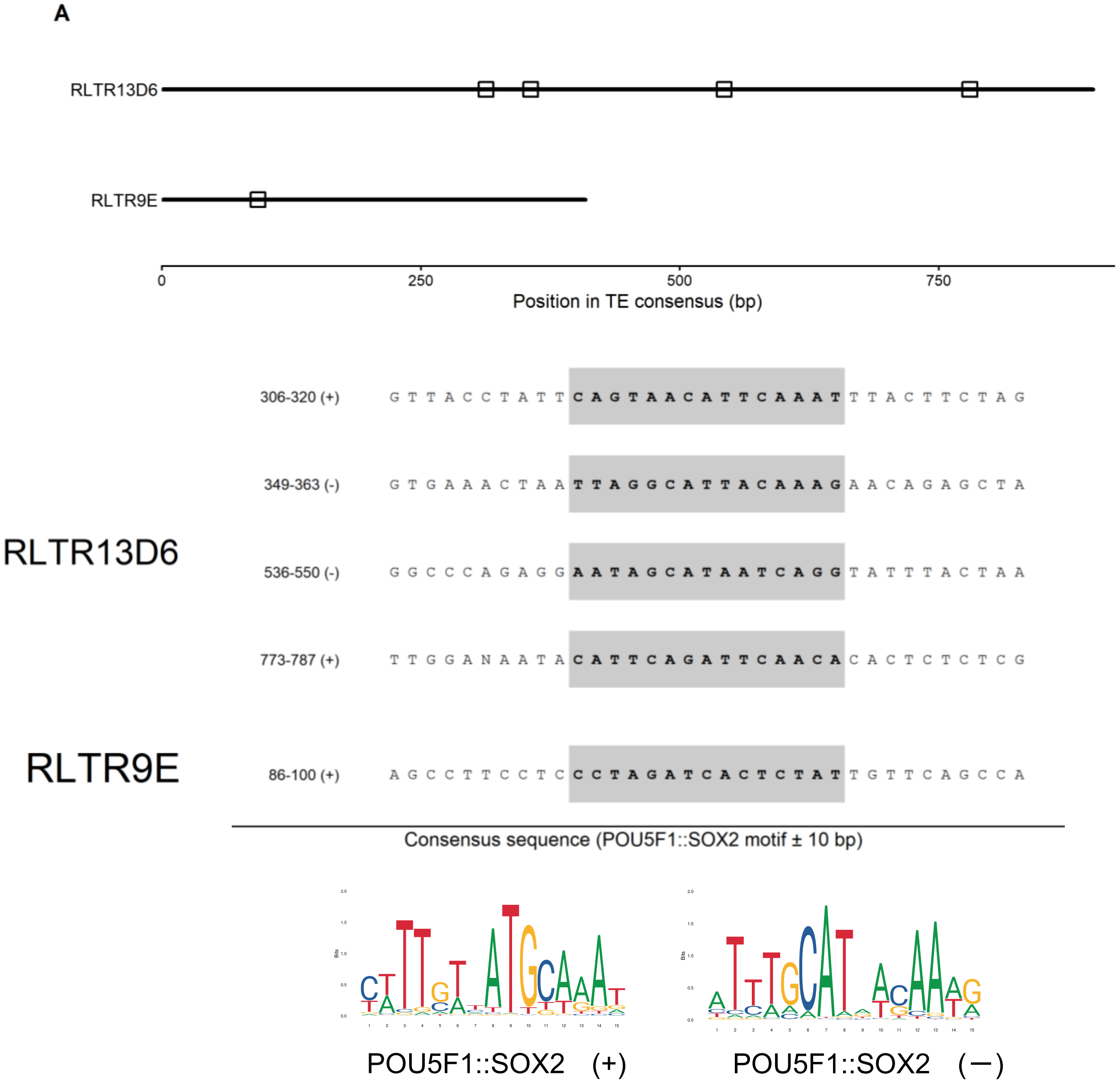


**Fig.S2 POU5F1::SOX2 motifs in the RLTR13D6 and RLTR9E consensus sequences.** Positions of POU5F1::SOX2 motif matches ($p < 0.001$ by FIMO) in the RLTR13D6 and RLTR9E consensus sequences. Sequences of the identified motif matches and the POU5F1::SOX2 motif logos are shown. The + and − in parenthesis indicate the strand orientation of each motif match.

Suzuki_Fig. S3

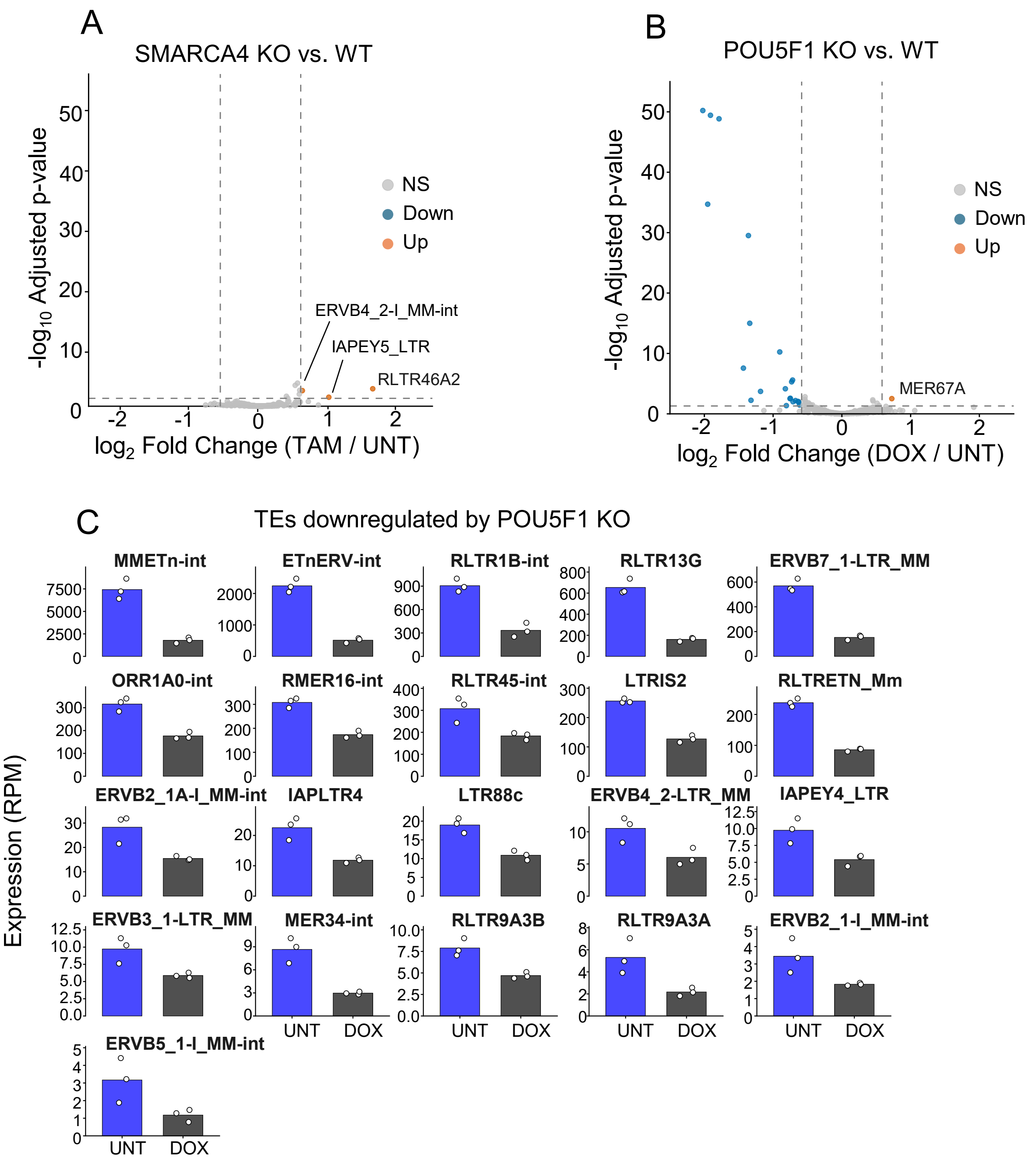


**Fig. S3. Changes in TE expression after SMARCA4 and POU5F1 depletion in mouse embryonic stem cells.**
**(A and B)** Volcano plots showing differential expression of TE subfamilies following SMARCA4 (A) and POU5F1 (B) depletion in mESCs. The x-axis represents the $\log_2$ fold change, and The y-axis represents the $-\log_{10}$ adjusted P-value. Significantly upregulated and downregulated TE subfamilies are shown in orange and blue, respectively (adjusted $P < 0.05$ and $|\log_2$ fold change$| > \log_2(1.5)$).
**(C)** Expression levels of TE subfamilies before (UNT) after (DOX) POU5F1. Expression levels were normalized as reads per million (RPM) based on the total counts of gene- and TE-mapped reads. Bars represent mean RPM values, and dots indicate individual technical replicates (n = 3 per condition).

Suzuki_Fig. S4

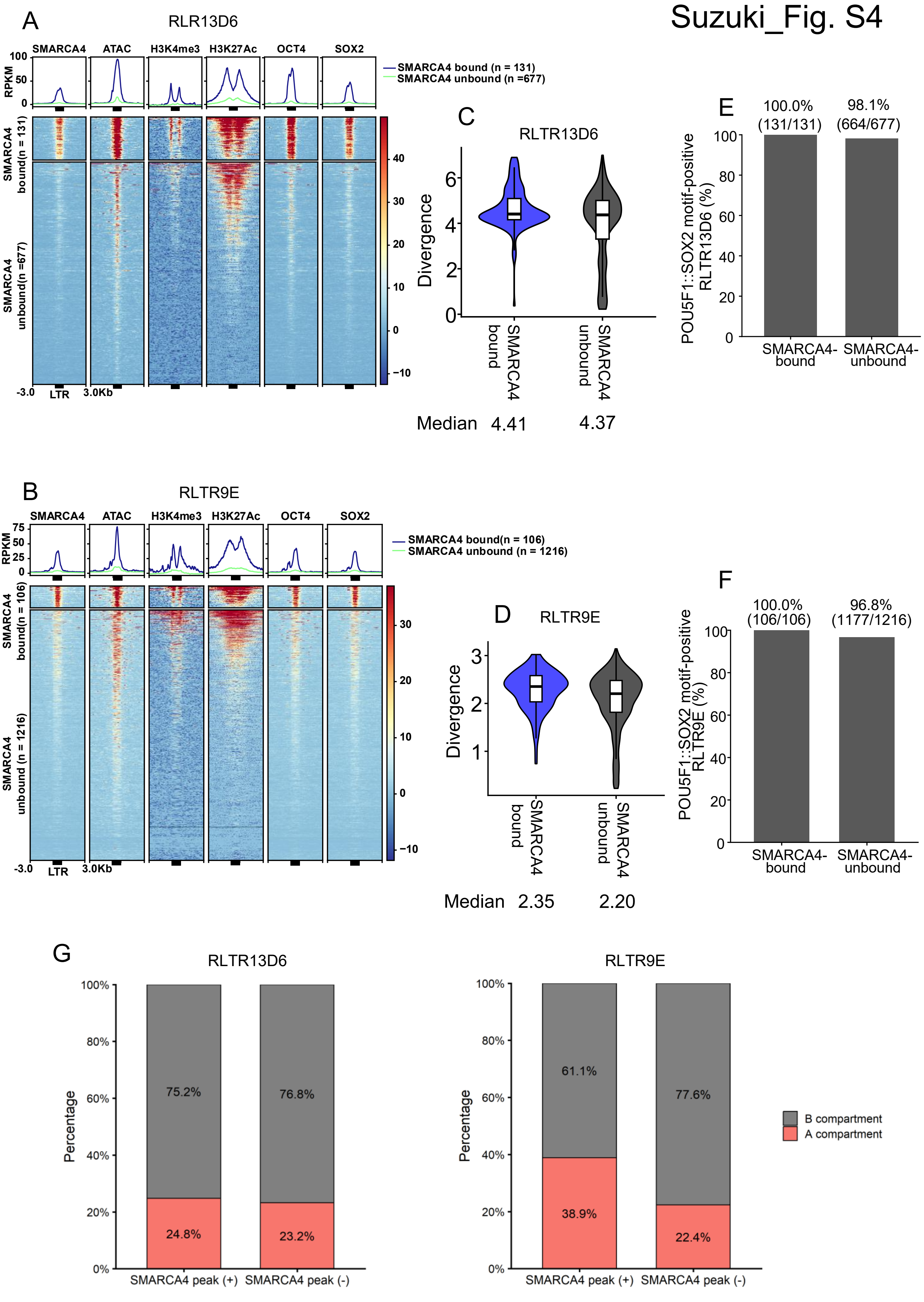


Figure legend is in the next page

**Fig.S4** Characteristics of SMARCA4-bound and unbound RLTR13D6 and RLTR9E copies. **(A and B)** SMARCA4, ATAC-seq, H3K4me3, H3K27ac, OCT4, and SOX2 signals at SMARCA4-bound and unbound RLTR13D6 **(A)** and RLTR9E **(B)** copies. Mapped reads are normalized as reads per kilobase per million mapped reads (RPKM) and shown as heatmaps and aggregated line plots. **(C and D)** Sequence divergence from the RepeatMasker consensus sequences of SMARCA4-bound and unbound RLTR13D6 **(C)** and RLTR9E **(D)** copies. **(E and F)** Fractions of SMARCA4-bound and unbound RLTR13D6 **(E)** and RLTR9E **(F)** copies containing POU5F1::SOX2 motif matches. **(G)** A/B compartment distributions of SMARCA4-bound and unbound RLTR13D6 and RLTR9E copies.

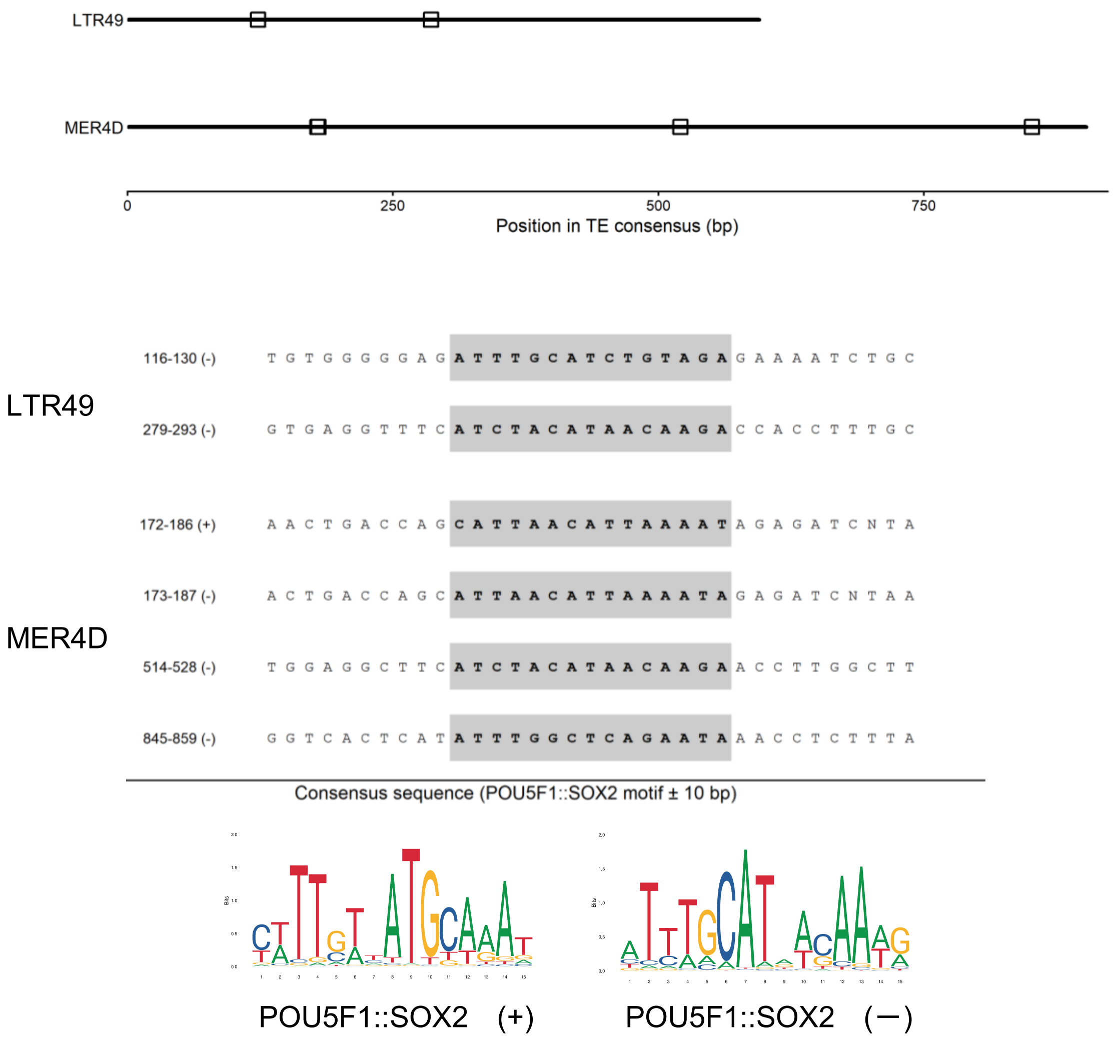


**Fig.S5 POU5F1::SOX2 motifs in LTR49 and MER4D consensus sequences.**
Positions of POU5F1::SOX2 motif matches (FIMO, $P < 0.001$) in the LTR49 and MER4D consensus sequences. Sequences of the identified motif matches and the POU5F1::SOX2 motif logos are shown. The + and − signs indicate the strand orientation of each motif match.
